# Assessing the relationship of ancient and modern populations

**DOI:** 10.1101/113779

**Authors:** Joshua G. Schraiber

## Abstract

Genetic material sequenced from ancient samples is revolutionizing our understanding of the recent evolutionary past. However, ancient DNA is often degraded, resulting in low coverage, error-prone sequencing. Several solutions exist to this problem, ranging from simple approach such as selecting a read at random for each site to more complicated approaches involving genotype likelihoods. In this work, we present a novel method for assessing the relationship of an ancient sample with a modern population while accounting for sequencing error and post-mortem damage by analyzing raw read from multiple ancient individuals simultaneously. We show that when analyzing SNP data, it is better to sequence more ancient samples to low coverage: two samples sequenced to 0.5x coverage provide better resolution than a single sample sequenced to 2x coverage. We also examined the power to detect whether an ancient sample is directly ancestral to a modern population, finding that with even a few high cover-age individuals, even ancient samples that are very slightly diverged from the modern population can be detected with ease. When we applied our approach to European samples, we found that no ancient samples represent direct ancestors of modern Europeans. We also found that, as shown previously, the most ancient Europeans appear to have had the smallest effective population sizes, indicating a role for agriculture in modern population growth.

## 1 Introduction

Ancient DNA (aDNA) is now ubiquitous in population genetics. Advances in DNA isolation [Dabney et al., 2013], library preparation [Meyer et al., 2012], bone sampling [Pinhasi et al., 2015], and sequence capture Haak et al. [2015] make it possible to obtain genome-wide data from hundreds of samples [Haak et al., 2015, Mathieson et al., 2015, Allentoft et al., 2015, Fu et al., 2016]. Analysis of these data can provide new insight into recent evolutionary processes which leave faint signatures in modern genomes, including natural selection [Schraiber et al., 2016, Jewett et al., 2016] and population replacement [Sjödin et al., 2014, Lazaridis et al., 2014].

One of the most powerful uses of ancient DNA is to assess the continuity of ancient and modern populations. In many cases, it is unclear whether populations that occupied an area in the past are the direct ancestors of the current inhabitants of that area. However, this can be next to impossible to assess using only modern genomes. Questions of population continuity and replacement have particular relevance for the spread of cultures and technology in humans [Lazaridis et al., 2016]. For instance, recent work showed that modern South Americans are descended from people associated Clovis culture that inhabited North America over 10,000 years ago, providing further evidence toward our understanding of the peopling of the Americas [Rasmussen et al., 2014].

Despite its utility in addressing difficult-to-answer questions in evolutionary biology, aDNA also has several limitations. Most strikingly, DNA decays rapidly following the death of an organism, resulting in highly fragmented, degraded starting material when sequencing [Sawyer et al., 2012]. Thus, ancient data is frequently sequenced to low coverage and has a significantly higher rate of misleadingly called nucleotides than modern samples. When working with diploid data, as in aDNA extracted from plants and animals, the low coverage prevents genotypes from being called called with confidence.

Several strategies are commonly used to address the low-coverage data. One of the most common approaches is to sample a random read from each covered site and use that as a haploid genotype call [Skoglund et al., 2012, Haak et al., 2015, Mathieson et al., 2015, Allentoft et al., 2015, Fu et al., 2016, Lazaridis et al., 2016]. Many common approaches to the analyses of ancient DNA, such as the usage of F-statistics [Green et al., 2010, Patterson et al., 2012], are designed with this kind of dataset in mind. As shown by Peter [2016], F-statistics can be interpreted as linear combinations of simpler summary statistics and can often be understood in terms of testing a tree-like structure relating populations. Nonetheless, despite the simplicity and appeal of this approach, it has several drawbacks. Primarily, it throws away reads from sites that are covered more than once, resulting in a potential loss of information from expensive, difficult-to-acquire data. These approach are also strongly impacted by sequencing error, post-mortem damage, and contamination.

On the other hand, several approaches exist to either work with genotype likelihoods or the raw read data. Genotype likelihoods are the probabilities of the read data at a site given each of the three possible diploid genotypes at that site. They can be used in calculation of population genetic statistics or likelihood functions to average over uncertainty in the genotype [Korneliussen et al., 2014]. However, many such approaches assume that genotype likelihoods are fixed by the SNP calling algorithm (although they may be recalibrated to account for aDNA-specific errors, as in **?**). However, with low coverage data, an increase in accuracy is expected if genotype likelihoods are co-estimated with other parameters of interest, due to the covariation between processes that influence read quality and genetic diversity, such as contamination.

A recent method that coestimates demographic parameters along with error and contamination rates by using genotype likelihoods showed that there can be significant power to assess the relationship of a single ancient sample to a modern population [Racimo et al., 2016]. Nonetheless, they found that for very low coverage data, inferences were not reliable. Thus, they were unable to apply their method to the large number of extremely low coverage (< 1x) genomes that are available. Moreover, they were unable to explore the tradeoffs that come with a limited budget: can we learn more by sequencing fewer individuals to high coverage, or more individuals at lower coverage?

Here, we develop a novel maximum likelihood approach for analyzing low coverage ancient DNA in relation to a modern population. We work directly with raw read data and explicitly model errors due to sequencing and port-mortem damage. Crucially, our approach incorporates data from multiple individuals that belong to the same ancient population, which we show substantially increases power and reduces error in parameter estimates. We then apply our new methodology to ancient human data, and show that we can perform accurate demographic inference even from very low coverage samples by analyzing them jointly.

## 2 Methods

### 2.1 Sampling alleles in ancient populations

We assume a scenario in which allele frequencies are known with high accuracy in a modern population. Suppose that an allele is known to be at frequency *x* ∈ (0, 1) in the modern population, and we wish to compute the probability of obtaining *k* copies of that allele in a sample of *n* (0 ≤ *k* ≤ *n*) chromosomes from an ancient population. Conditioning on the frequency of the allele in the modern population minimizes the impact of ascertainment, and allows this approach to be used for SNP capture data.

To calculate the sampling probability, we assume a simple demographic model in which the ancient individual belongs to a population that split off from the modern population *τ*_1_ generations ago, and subsequently existed as an isolated population for *τ*_2_ generations. Further, we assume that the modern population has effective size 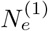 and that the ancient population has effective size 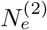, and measure time in diffusion units,inline equation 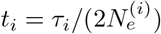
If we know the conditional probability that an allele is at frequency *y* in the ancient sample, given that it is at frequency *x*, denoted *f*(*y*; *x, t*_1_, *t*_2_), then the sampling probability is simply an integral,

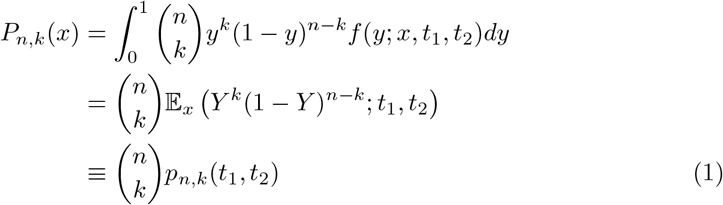

Thus, we must compute the binomial moments of the allele frequency distribution in the ancient population. In the Appendix, we show that this can be computed using matrix exponentiation,

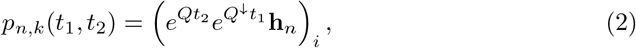

where (**v**)_*i*_ indicates the *i*th element of the vector **v**, **h**_*n*_ = ((1*−x*)^*n*^, x(1*−x*)^*n−*1^, *…, x^n^*)^*T*^ and *Q* and *Q^↓^* are the sparse matrices

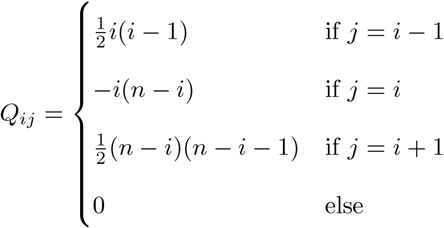

and

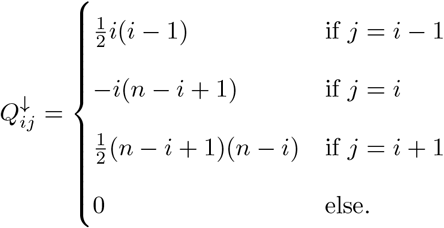

This result has an interesting interpretation: the matrix *Q^↓^* can be thought of as evolving the allele frequencies back in time from the modern population to the common ancestor of the ancient and modern populations, while *Q* evolves the allele frequencies forward in time from the common ancestor to the ancient population (Fig 1).

**Figure 1:**
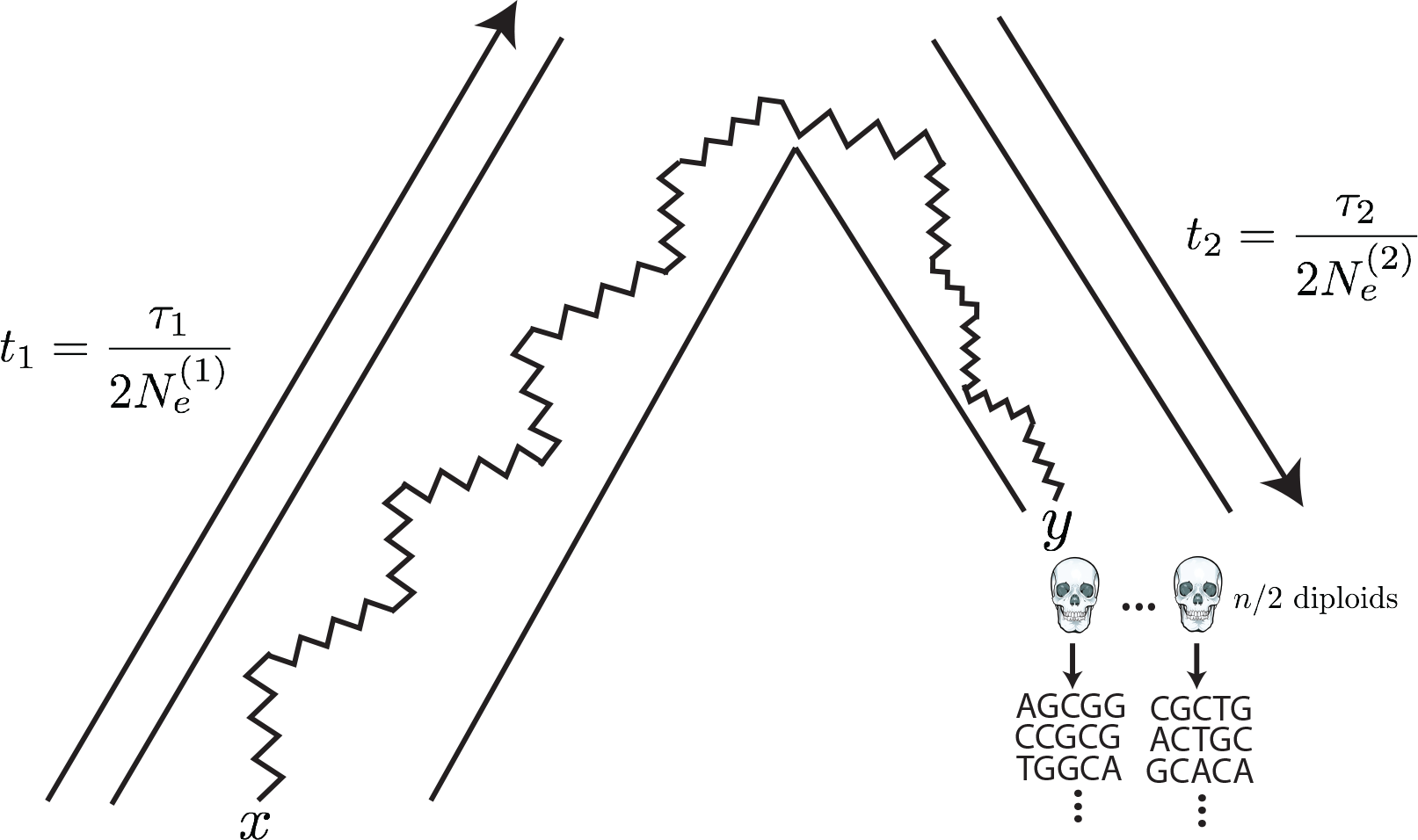
The generative model. Alleles are found at frequency *x* in the modern population and are at frequency *y* in the ancient population. The modern population has effective size 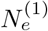 and has evolved for *Τ*_1_ generations since the common ancestor of the modern and ancient populations, while the ancient population is of size 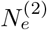 and has evolved for *Τ*_2_ generations. Ancient diploid samples are taken and sequenced to possibly low coverage, with errors. Arrows indicate that the sampling probability can be calculated by evolving alleles *backward* in time from the modern population and then forward in time to the ancient population.

Because of the fragmentation and degradation of DNA that is inherent in obtaining sequence data from ancient individuals, it is difficult to obtain the high coverage data necessary to make high quality genotype calls from ancient individuals. To address this, we instead work directly with raw read data, and average over all the possible genotypes weighted by their probability of producing the data. Specifically, we follow Nielsen et al. [2012] in modeling the probability of the read data in the ancient population, given the allele frequency at site *l* as

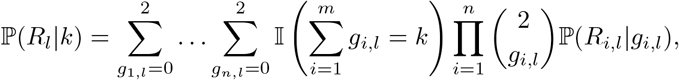

where *R_i,l_* = (*a_i,l_, d_i,l_*) are the counts of ancestral and derived reads in individual *i* at site *l*, *g_i,l_* ∈ {0, 1, 2} indicates the possible genotype of individual *i* at site *l* (i.e. 0 = homozygous ancestral, 1 = heterozygous, 2 = homozygous derived), and ℙ(*R_i,l_|g_i,l_*) is the probability of the read data at site *l* for individual *i*, assuming that the individual truly has genotype *g_i,l_*. We use a binomial sampling with error model, in which the probability that a truly derived site appears ancestral (and vice versa) is given by *ϵ*. We emphasize that the parameter *ϵ* will capture both sequencing error as well as post-mortem damage (c.f. Racimo et al. [2016] who found that adding an additional parameter to specifically model post-mortem damage does not improve inferences). Thus,

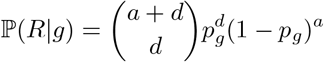

with

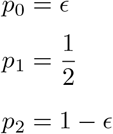

Combining these two aspects together by summing over possible allele frequencies weighted by their probabilities, we obtain our likelihood of the ancient data,

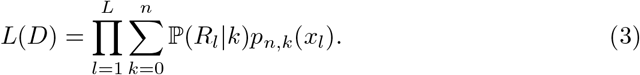

## 3 Results

### 3.1 Impact of coverage and number of samples on inferences

To explore the tradeoff of sequencing more individuals at lower depth compared to fewer individuals at higher coverage, we performed simulations using msprime [Kelleher et al., 2016] combined with custom scripts to simulate error and low coverage data. Briefly, we assumed a Poisson distribution of reads at every site with mean given by the coverage, and then simulated reads by drawing from the binomial distribution described in the Methods.

First, we examined the impact of coverage and number of samples on the ability to recover the drift times in the modern and the ancient populations. Figure 2 shows results for data simulated with *t*_1_ = 0.02 and *t*_2_ = 0.05, corresponding to an ancient individual who died 300 generations ago from population of effective size 1000. The populations split 400 generations ago, and the modern population has an effective size of 10000. We simulated approximately 180000 SNPs by simulating 100000 500 base pair fragments. Inferences of *t*_1_ can be relatively accurate even with only one low coverage ancient sample (Figure 2A). However, inferences of *t*_2_ benefit much more from increasing the number of ancient samples, as opposed to coverage (Figure 2B). In particular, two individuals sequenced to 0.5x coverage have a much lower error than a single individual sequenced to 2x coverage. To explore this effect further, we derived the sampling probability of alleles covered by exactly one sequencing read (see Appendix). We found that sites covered only once have no information about *t*_2_, suggesting that evidence of heterozygosity is very important for inferences about *t*_2_.

**Figure 2:**
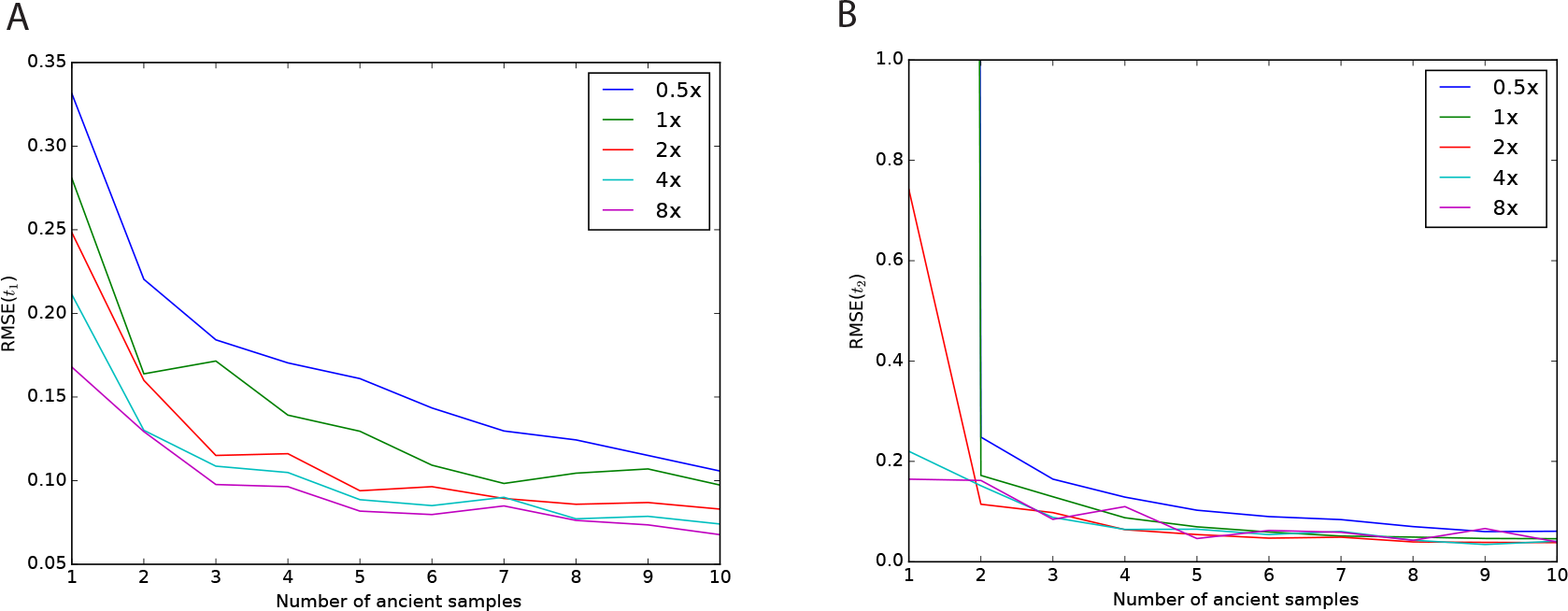
Impact of sampling scheme on parameter estimation error. In each panel, the *x* axis represents the number of simulated ancient samples, while the *y* axis shows the relative root mean square error for each parameter. Each different line corresponds to individuals sequenced to different depth of coverage. Panel A shows results for *t*_1_ while panel B shows results for *t*_2_. Simulated parameters are *t*_1_ = 0.02 and *t*_2_ = 0.05.

We next examined the impact of coverage and sampling on the power to reject the hypothesis that the ancient individuals came from a population that is directly ancestral to the modern population. We analyzed both low coverage (0.5x) and higher coverage (4x) datasets consisting of 1 (for both low and high coverage samples) or 5 individuals (only for low coverage). We simulated data with parameters identical to the previous experiment, except we now examined the impact of varying the age of the ancient sample from 0 generations ago through to the split time with the modern population. We then performed a likelihood ratio test comparing the null model of continuity, in which *t*_2_ = 0, to a model in which the ancient population is not continuous. Figure 3 shows the power of the likelihood ratio test. For a single individual sequenced to low coverage, we see that the test only has power for very recently sampled ancient individuals (i.e. samples that are highly diverged from the modern population). However, the power increases dramatically as the number of individuals or the coverage per individual is increased; sequencing 5 individuals to 0.5x coverage results in essentially perfect power to reject continuity. Nonetheless, for samples that are very close to the divergence time, it will be difficult to determine if they are ancestral to the modern population or not, because differentiation is incomplete.

**Figure 3:**
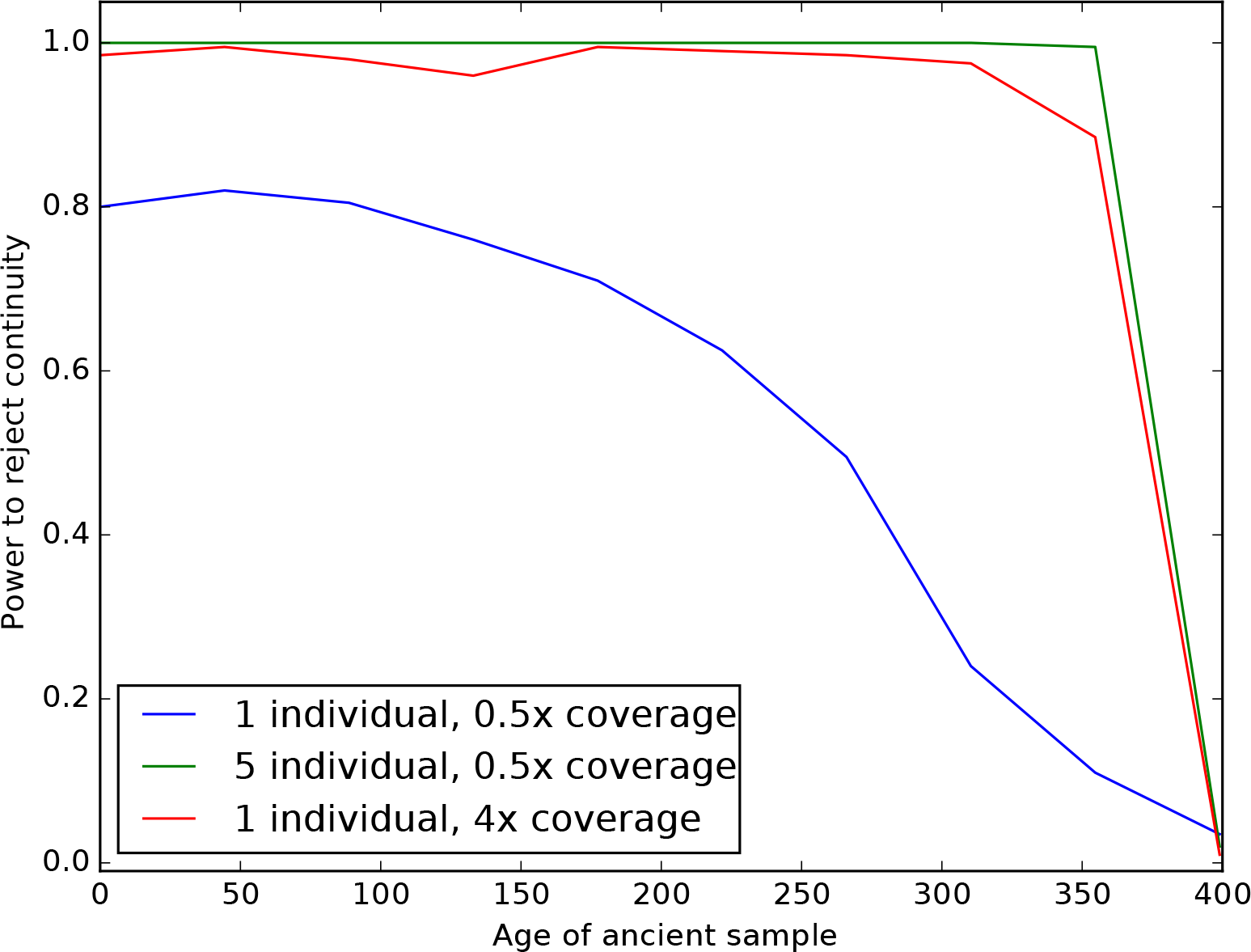
Impact of sampling scheme on rejecting population continuity. The *x* axis represents the age of the ancient sample in generations, with 0 indicating a modern sample and 400 indicating a sample from exactly at the split time 400 generations ago. The *y* axis shows the proportion of simulations in which we rejected the null hypothesis of population continuity. Each line shows different sampling schemes, as explained in the legend.

### 3.2 Impact of admixture

We examined two possible violations of the model to assess their impact on inference. In many situations, there may have been secondary contact between the population from which the ancient sample is derived and the modern population used as a reference. We performed simulations of this situation by modifying the previous simulations to include subsequent admixture from the ancient population to the modern population 200 generations ago (NB: this admixture occurred *more recently* than the ancient sample). In Figure 4, we show the results for admixture proportions ranging from 0 to 50%. Counterintuitively, estimates of *t*_1_ initially *decrease* before again increasing. This is likely a result of the increased heterozygosity caused by admixture, which acts to artificially inflate the effective size of the modern population, and thus decrease *t*_1_. As expected, *t*_2_ is estimated to be smaller the more admixture there is; indeed, for an admixture rate of 100%, the modern and ancient samples are continuous. The impact on *t*_2_ appears to be linear, and is well approximated by (1 *− f*) *t*_2_ if the admixture fraction is *f*.

**Figure 4:**
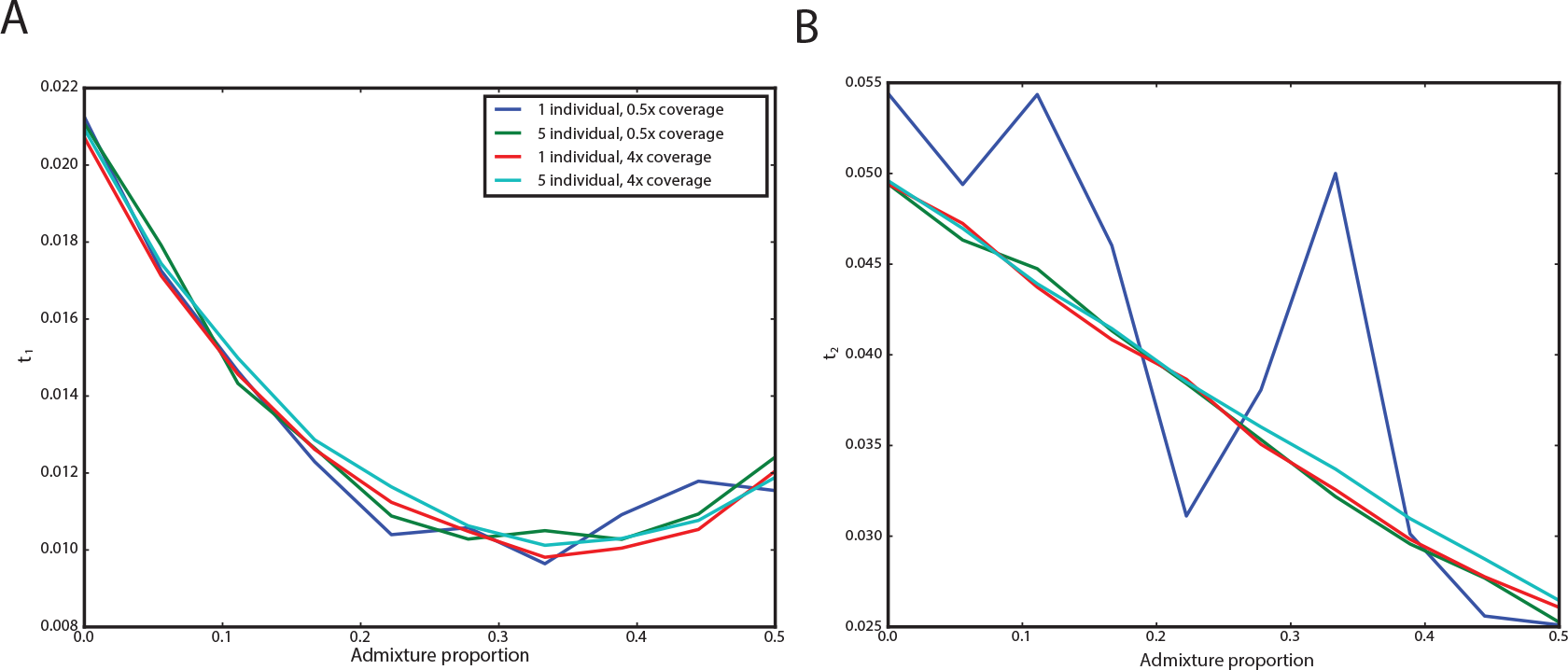
Impact of admixture from the ancient population on inferred parameters. The *x* axis shows the admixture proportion and the *y* axis shows the average parameter estimate across simulations. Each line corresponds to a different sampling strategy, as indicated in the legend. Panel A shows results for *t*_1_ and Panel B shows results for *t*_2_.

In other situations, there may be admixture from an unsampled “ghost” population into the modern population. If the ghost admixture is of a high enough proportion, it is likely to cause a sample that is in fact a member of a directly ancestral population to not appear to be ancestral. We explored this situation by augmenting our simulations in which the ancient sample is continuous with an outgroup population diverged from the modern population 0.04 time units ago (corresponding to 800 generations ago) and contributed genes to the modern population 0.01 time units ago (corresponding to 200 generations ago). We then assessed the impact on rejecting continuity using the likelihood ratio test (Figure 5). As expected, we see that low-power sampling strategies (such as a single individual sequenced to low coverage) are relatively unimpacted by ghost admixture. However, even relatively powerful sampling strategies can be robust to ghost admixture up to approximately 10%.

**Figure 5.**
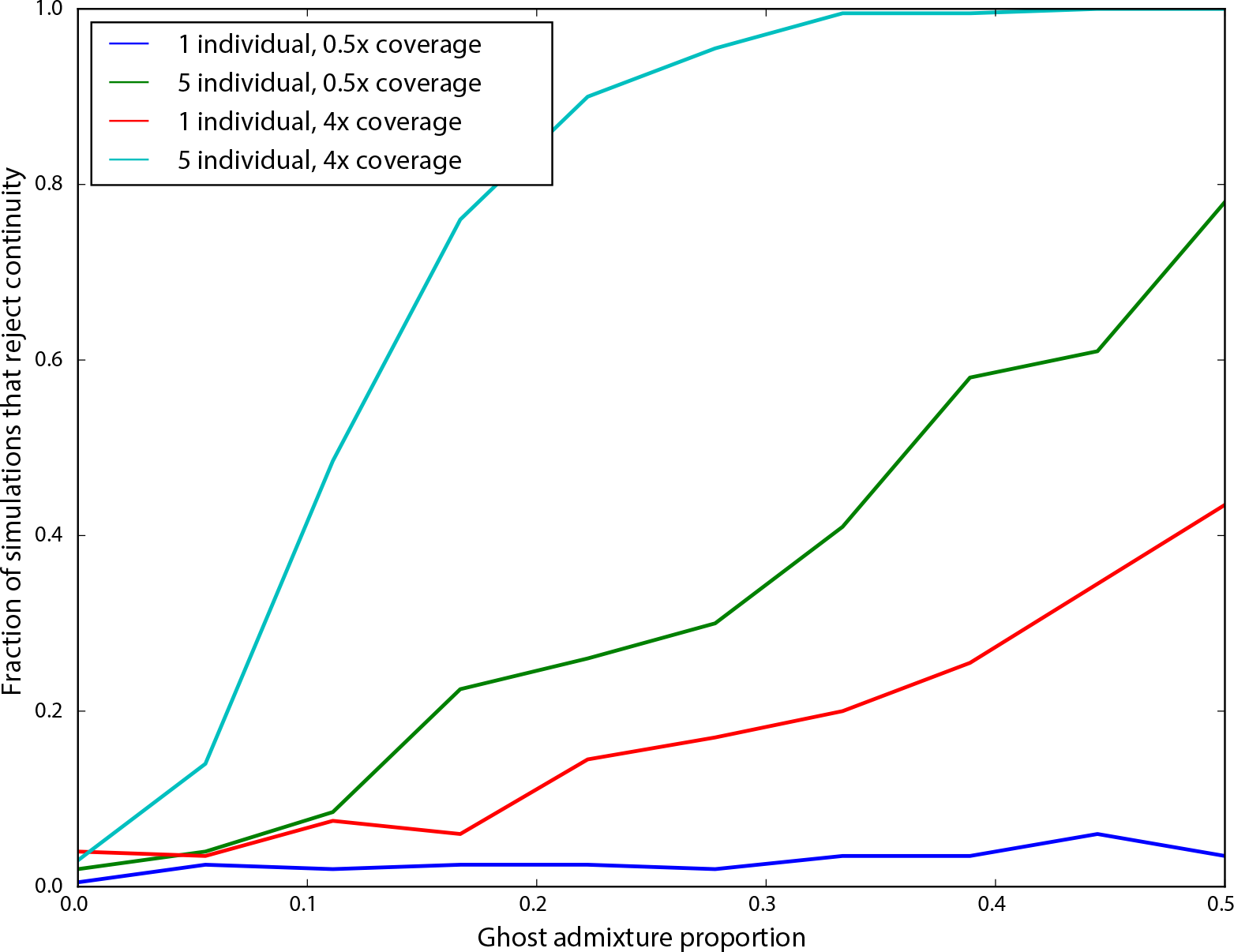
Impact of ghost admixture on falsely rejecting continuity. The *x* axis shows the admixture proportion from the ghost population, and the *y* axis shows the fraction of simulations in which continuity was rejected. Each line corresponds to a different sampling strategy, as indicated in the legend.

### 3.3 Application to ancient humans

We applied our approach to ancient human data from Mathieson et al. [2015], which is primarily derived from a SNP capture approach that targeted 1.2 million SNPs. Based on sampling location and associated archeological materials, the individuals were grouped into *a priori* panels, which we used to specify population membership when analyzing individuals together. We analyzed all samples for their relationship to the CEU individuals from the 1000 Genomes Project [Consortium, 2015]. Based on our results that suggested that extremely low coverage samples would yield unreliable estimates, we excluded panels that are composed of only a single individual sequenced to less than 2x coverage.

We computed maximum likelihood estimates of *t*_1_ and *t*_2_ for individuals as grouped into populations (Figure 6A; Table 1). We observe that *t*_2_ is significantly greater than 0 for all populations. Thus, none of these populations are consistent with directly making up a large proportion of the ancestry of modern CEU individuals. Strikingly, we see that *t*_2_ ≫ *t*_1_, despite the fact that the ancient samples must have existed for fewer generations since the population split than the modern samples. This suggests that all of the ancient populations are characterized by extremely small effective population sizes.

**Figure 6:**
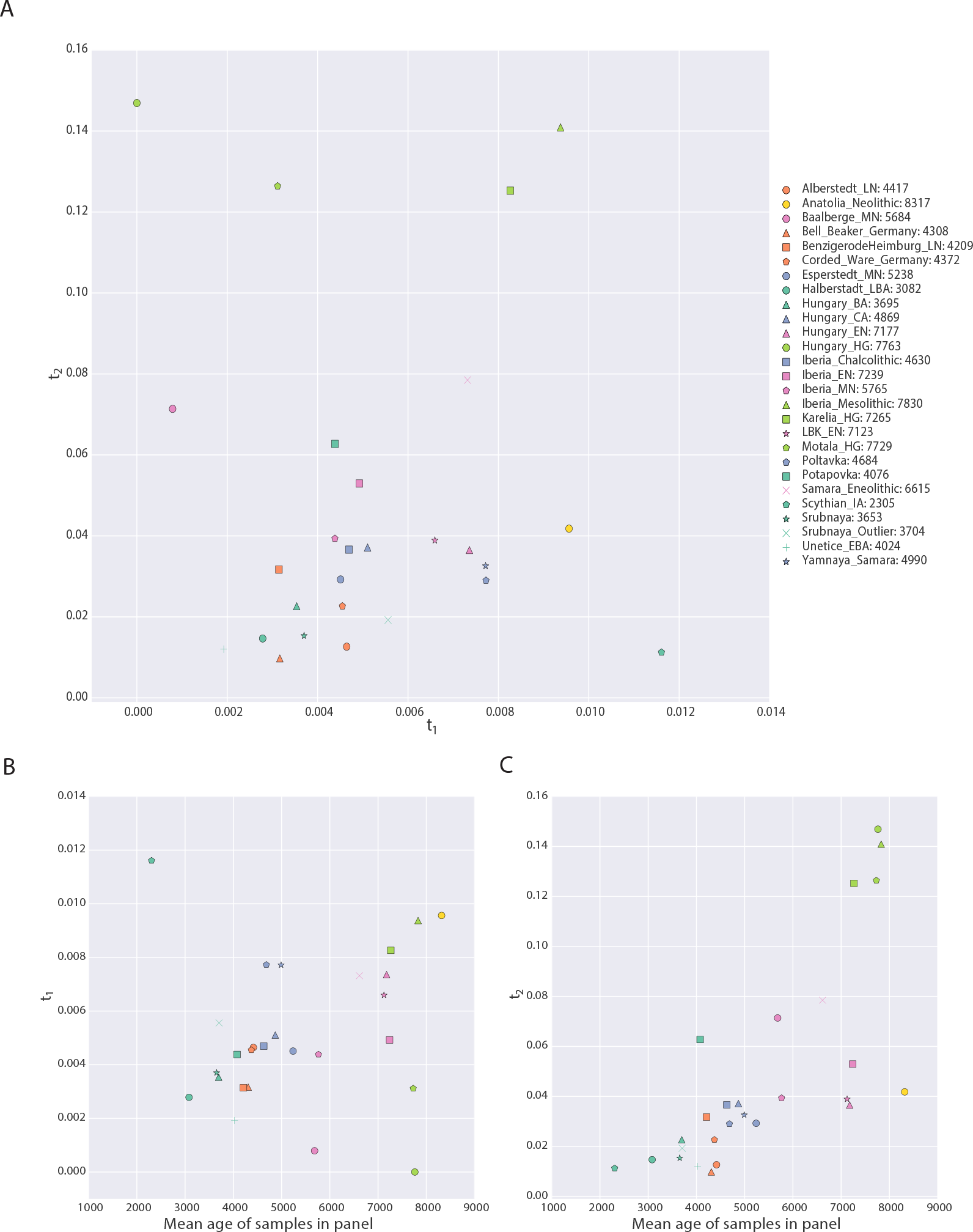
Parameters of the model inferred from ancient West Eurasian samples. Panel A shows *t*_1_ on the x-axis and *t*_2_ on the y-axis, with each point corresponding to a population as indicated in the legend. Numbers in the legend correspond to the mean date of all samples in the population. Panels B and C show scatterplots of the mean age of the samples in the population (*x*-axis) against *t*_1_ and *t*_2_, respectively. Points are described by the same legend as Panel A.

**Table 1:**
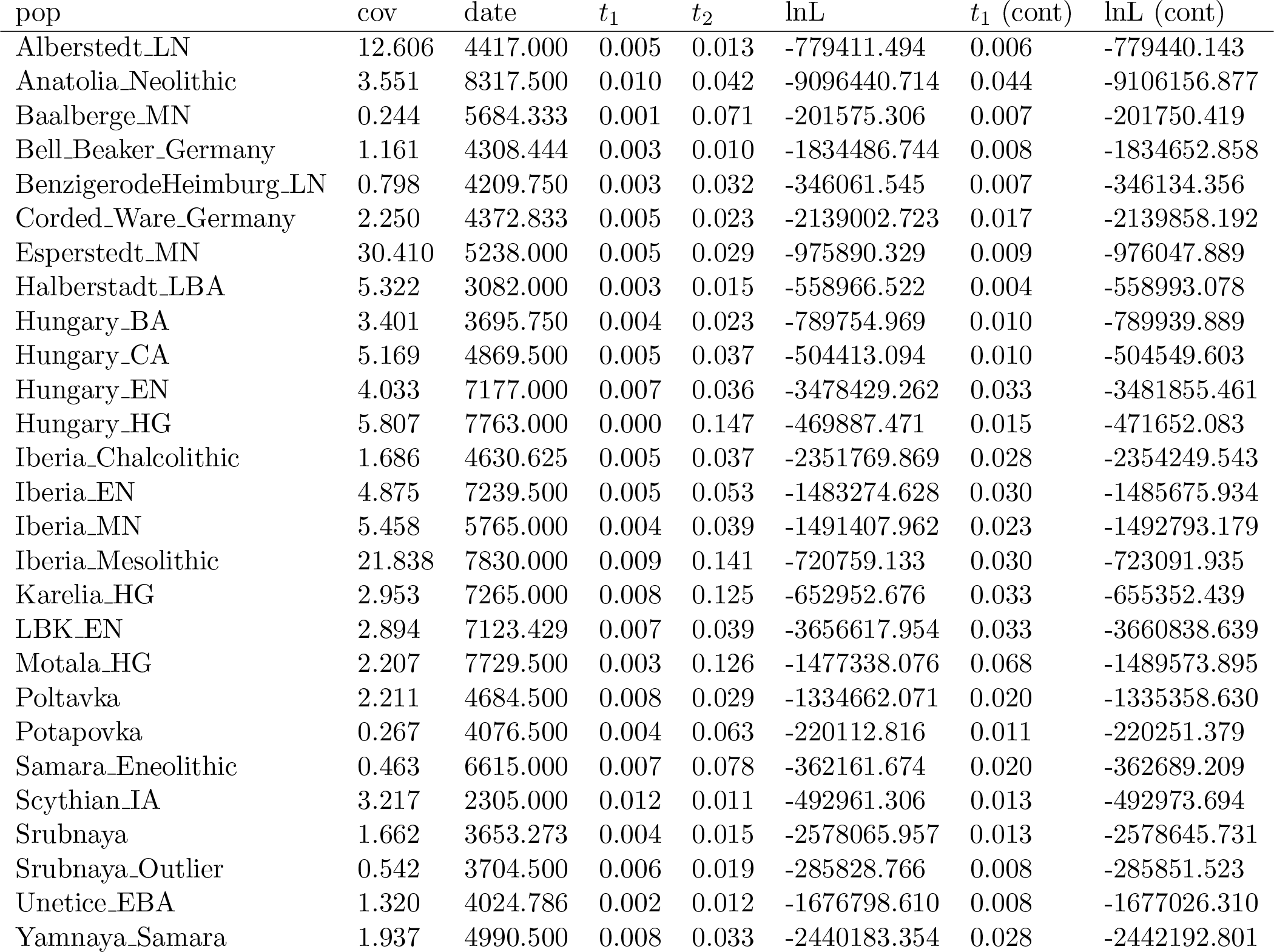
Details of populations included in analysis. “pop” is population name, “cov” is mean coverage of individuals in the population, “date” is mean date of individuals in the population, “*t*_1_” is the maximum likelihood estimate of *t*_1_ in the full model, “*t*_2_” is the maximum likelihood estimate of *t*_2_ in the full model, “LnL” is the maximum likelihood value in the full model, “*t*_1_ (cont)” is the maximum likelihood estimate of *t*_1_ in the model where *t*_2_ = 0, “LnL” is the maximum likelihood value in the model where *t*_2_ = 0.

We further explored the relationship between the dates of the ancient samples and the parameters of the model by plotting *t*_1_ and *t*_2_ against the mean sample date of all samples in that population (Figure 6B, C). We expected to find that *t*_1_ correlated with sample age, under the assumption that samples were members of relatively short-lived populations that diverged from the “main-stem” of CEU ancestry. Instead, we see no correlation between *t*_1_ and sample time, suggesting that the relationship of these populations to the CEU is complicated and not summarized well by the age of the samples. On the other hand, we see a strong positive correlation between *t*_2_ and sampling time (*p <* 1 *×* 10^*−*4^). Because *t*_2_ is a compound parameter, it is difficult to directly interpret this relationship. However, it is consistent with the most ancient samples belonging to populations with the smallest effective sizes, consistent with previous observations [Skoglund et al., 2014].

Finally, we examined the impact of grouping individuals into populations in real data. We see that estimates of *t*_1_ for low coverage samples are typically lower when analyzed individually than when pooled with other individuals of the same panel (Figure 7A), suggesting a slightly downward bias in estimating *t*_1_ for low coverage samples. On the other hand, there is substantial bias toward overestimating *t*_2_ when analyzing samples individually, particularly for very low coverage samples (Figure 7B). This again shows that for estimates that rely on heterozygosity in ancient populations, pooling many low coverage individuals can significantly improve estimates.

**Figure 7:**
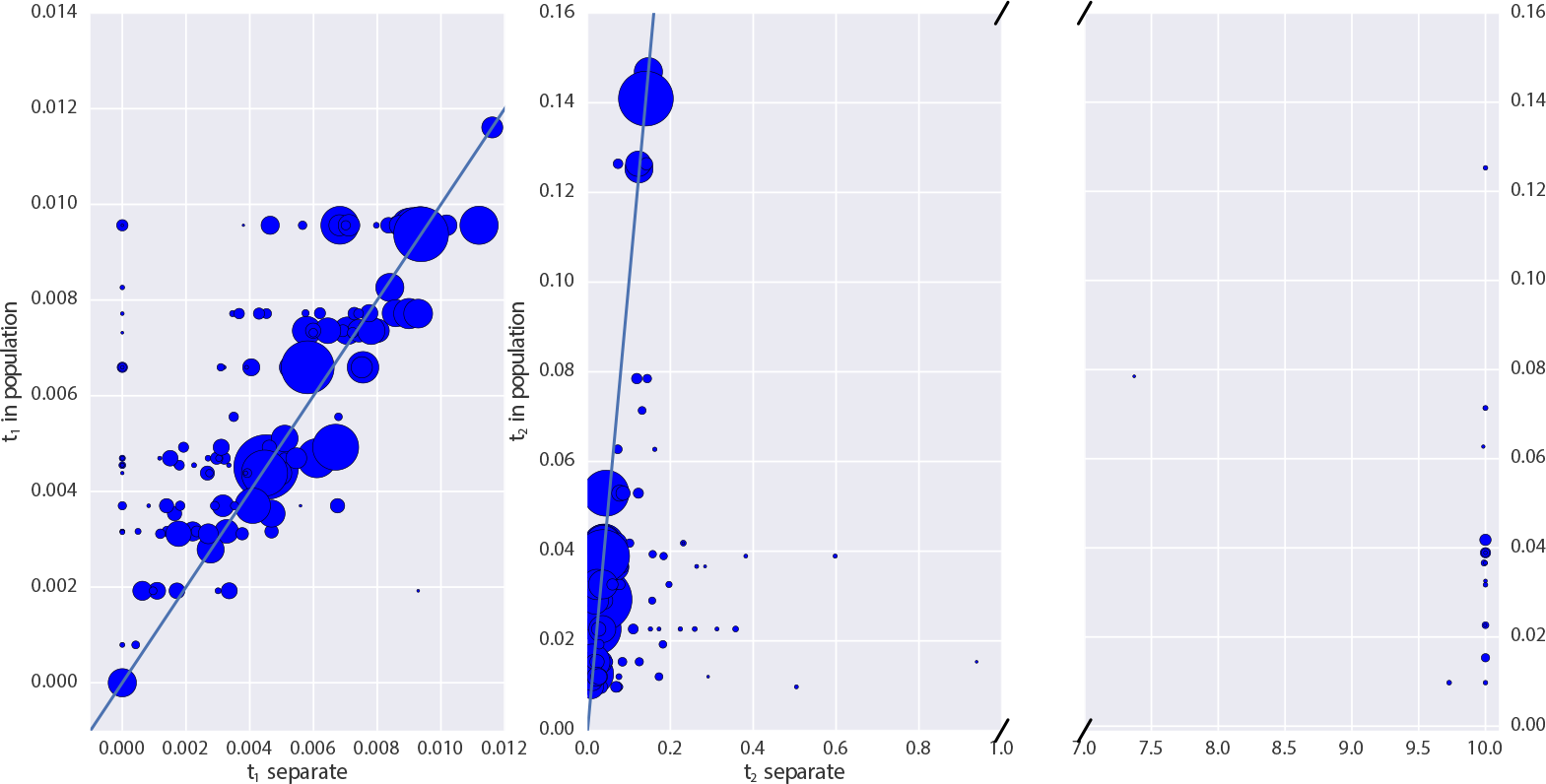
Impact of pooling individuals into populations when estimating model parameters from real data. In both panels, the x-axis indicates the parameter estimate when individuals are analyzed separately, while the y-axis indicates the parameter estimate when individuals are grouped into populations. Size of points is proportional to the coverage of each individual. Panel A reports the impact on estimation of *t*_1_, while Panel B reports the impact on *t*_2_. Note that Panel B has a broken x-axis. Solid lines in each figure indicate *y* = *x*.

## 4 Discussion

Ancient DNA (aDNA) presents unique opportunities to enhance our understanding of demography and selection in recent history. However, it also comes equipped with several challenges, due to postmortem DNA damage [Sawyer et al., 2012]. Several strategies have been developed to deal with the low quality of aDNA data, from relatively simple options like sampling a read at random at every site [Green et al., 2010] to more complicated methods making use of genotype likelihoods [Racimo et al., 2016]. Here, we presented a novel maximum likelihood approach for making inferences about how ancient populations are related to modern populations by analyzing read counts from multiple ancient individuals and explicitly modeling relationship between the two populations. We explicitly condition on the allele frequency in a modern population; thus, our method is robust to ascertainment in modern samples and can be used with SNP capture data. Using this approach, we examined some aspects of sampling strategy for aDNA analysis and we applied our approach to ancient humans.

We found that sequencing many individuals from an ancient population to low coverage (.5-1x) can be a significantly more cost effective strategy than sequencing fewer individuals to relatively high coverage. For instance, we saw from simulations that far more accurate estimates of the drift time in an ancient population can be obtained by pooling 2 individuals at 0.5x coverage than by sequencing a single individual to 2x coverage (Figure 2). We saw this replicated in our analysis of the real data: low coverage individuals showed a significant amount of variation and bias in estimating the model parameters that was substantially reduced when individuals were analyzed jointly in a population (Figure 7). To explore this further, we showed that sites sequenced to 1x coverage in a single individual retain no information about the drift time in the ancient population. This can be intuitively understood because the drift time in the ancient population is strongly related the amount of heterozygosity in the ancient population: an ancient population with a longer drift time will have lower heterozygosity at sites shared with a modern population. When a site is only sequenced once in a single indi-vidual, there is no information about the heterozygosity of that site. We also observed a pronounced upward bias in estimates of the drift time in the ancient population from low coverage samples. We speculate that this is due to the presence of few sites covered more than once being likely to be homozygous, thus deflating the estimate of heterozygosity in the ancient population. Thus, for analysis of SNP data, we recommend that aDNA sampling be conduced to maximize the number of individuals from each ancient population that can be sequenced to ∼1x, rather than attempting to sequence fewer individuals to high coverage.

When we looked at the impact of model misspecification, we saw several important patterns. First, the influence of admixture from the ancient population on inferences of *t*_2_ is approximately linear, suggesting that if there are estimates of the amount of admixture between the modern and ancient population, a bias-corrected estimate of *t*_2_ could be produced (Figure 4B). The impact on inference of *t*_1_ is more complicated: admixture actually *reduces* estimates of *t*_1_ (Figure 4A). This is likely because admixture increases the heterozygosity in the modern population, thus causing the amount of drift time to seem reduced. In both cases, the bias is not impacted by details of sampling strategy, although the variance of estimates is highly in a way consistent with Figure 2.

Of particular interest in many studies of ancient populations is the question of direct ancestry: are the ancient samples members of a population that contributed substantially to a modern population? We emphasize that this does not mean that the particular samples were direct ancestors of any modern individuals; indeed, this is exceedingly unlikely for old samples [Rohde et al., 2004, Chang, 1999, Baird et al., 2003, Donnelly, 1983]. Instead, we are asking whether an ancient sample was a member of a population that is directly continuous with a modern population. Several methods have been proposed to test this question, but thus far they have been limited to many individuals sequenced at a single locus [Sjödin et al., 2014] or to a single individual with genome-wide data [Rasmussen et al., 2014]. Our approach provides a rigorous, maximum likelihood framework for testing questions of population continuity using multiple low coverage ancient samples. We saw from simulations (Figure 3) that data from single, low coverage individuals result in very little power to reject the null hypothesis of continuity unless the ancient sample is very recent (i.e. it has been diverged from the modern population for a long time). Nonetheless, when low coverage individuals are pooled together, or a single high coverage individual is used, there is substantial power to reject continuity for all but the most ancient samples (i.e. samples dating from very near the population split time).

Because many modern populations may have experienced admixture from unsampled “ghost” populations, we also performed simulations to test the impact of ghost admixture on the probability of falsely rejecting continuity. We find that single ancient samples do not provide sufficient power to reject continuity even for high levels of ghost admixture, while increasingly powerful sampling schemes, adding more individuals or higher coverage per individual, reject continuity at higher rates. However, in these situations, whether we regard rejection of continuity as a false or true discovery is somewhat subjective: how much admixture from an outside population is required before considering a population to not be directly ancestral? In future work it will be extremely important to estimate the “maximum contribution” of the population an ancient sample comes from (c.f Sjödin et al. [2014].

To gain new insights from empirical data, we applied our approach to ancient samples throughout Europe. Notably, we rejected continuity for all populations that we analyzed. This is unsurprising, given that European history is extremely complicated and has been shaped by many periods of admixture [Lazaridis et al., 2014, Haak et al., 2015, Lazaridis et al., 2016]. Thus, modern Europeans have experienced many periods of “ghost” admixture (relative to any particular ancient sample). Nonetheless, our results show that none of these populations are even particularly close to directly ancestral, as our simulations have shown that rejection of continuity is robust to low levels of ghost admixture.

Secondly, we observed that the drift time in the ancient population was much larger than the drift time in the modern population. Assuming that the ancient sample were a contemporary sample, the ratio *t*_1_/*t*_2_ is an estimator of the ratio inline equation 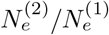; in fact, because the ancient sample existed for fewer generations since the common ancestor of the ancient and modern populations, *t*_1_/*t*_2_ acts as an upper bound oninline equation 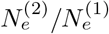. Moreover, this is unlikely to be due to unmodeled error in the ancient samples: error would be expected increase the heterozygosity in the ancient sample, and thus *decrease* our estimates of *t*_2_. Thus, we find strong support for the observation that ancient Europeans were often members of small, isolated populations [Skoglund et al., 2014]. We interpret these these two results together as suggestive that many ancient samples found thus far in Europe were members of small populations that ultimately went locally extinct.

We further examined the effective sizes of ancient populations through time by looking for a correlation between the age of the ancient populations and the drift time leading to them (Figure 6C). We saw a strong positive correlation, and although this drift time is a compound parameter, which complicates interpretations, it appears that the oldest Europeans were members of the smallest populations, and that effective population size has grown through time as agriculture spread through Europe.

We anticipate the further development of methods that explicitly account for differential drift times in ancient and modern samples will become important as aDNA research becomes even more integrating into population genomics. This is because many common summary methods, such as the use of Structure [Pritchard et al., 2000] and Admixture [Alexander et al., 2009], are sensitive to differential amounts of drift between populations [Falush et al., 2016]. As we’ve shown in ancient Europeans, ancient samples tend to come from isolated subpopulations with a large amount of drift, thus confounding such summary approaches. Moreover, standard population genetics theory shows that allele frequencies are expected to be deterministically lower in ancient samples, even if they are direct ancestors of a modern population. Intuitively, this arises because the alleles must have arisen at some point from new mutations, and thus were at lower frequencies in the past. A potentially fruitful avenue to combine these approaches moving forward may be to separate regions of the genome based on ancestry components, and assess the ancestry of ancient samples relative to specific ancestry components, rather than to genomes as a whole.

Our current approach leaves several avenues for improvement. We use a relatively simple error model that wraps up both post-mortem damage and sequencing error into a single parameter. While Racimo et al. [2016] shows that adding an additional parameter for PMD-related error does not significantly change results, the recent work of ? shows that building robust error models is challenging and essential to estimating heterozygosity properly. Although our method is robust to non-constant demography because we consider only alleles that are segregating in both the modern and the ancient population, we are losing information by not modeling new mutations that arise in the ancient population. Similarly, we only consider a single ancient population at a time, albeit with multiple samples. Ideally, ancient samples would be embedded in complex demographic models that include admixture, detailing their relationships to each other and to modern populations [Patterson et al., 2012, Lipson and Reich, 2017]. However, inference of such complex models is difficult, and though there has been some progress in simplified cases [Lipson et al., 2014, Pickrell and Pritchard, 2012], it remains an open problem due to the difficult of simultaneously inferring a non-tree-like topology along with demographic parameters. Software such as momi [Kamm et al., 2016] that can compute the likelihood of SNP data in an admixture graph may be able to be used to integrate over genotype uncertainty in larger settings than considered here.

## 5 Appendix

### 5.1 Computing allele frequency moments in the ancient population

We wish to compute moments of the form

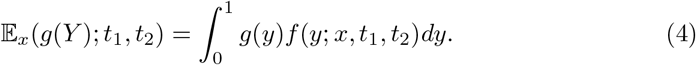

To do so, we make use of several results from diffusion theory. To ensure that this paper is self contained, we briefly review those results here. The interested reader may find much of this material covered in Ewens [2012], Karlin and Taylor [1981]. Several similar calculations can be found in Griffiths [2003].

Let the probability of an allele going from frequency *x* to frequency *y* in *Τ* generations in a population of size *N_e_* be *f* (*x, y*; *t*), where *t* = *Τ /*(2*N_e_*). Under a wide variety of models, the change in allele frequencies through time is well approximated by the Wright-Fisher diffusion, which is characterized by its generator,

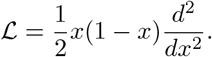

The generator of a diffusion process is useful, because it can be used to define a differential equation for the moments of that process,

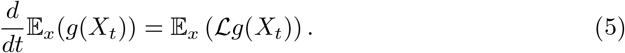

We will require the *speed measure* of the Wright-Fisher diffusion, *m*(*x*) = *x^−^*^1^(1 *− x*)^*−*1^, which essentially describes how slow a diffusion at position *x* is “moving” compared to a Brownian motion at position *x*. Note that all diffusions are reversible with respect to their speed measures, i.e

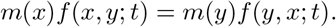

We additionally require the probability of loss, i.e. the probability that the allele currently at frequency *x* is ultimately lost from the population. This is

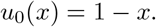

Note that it is possible to condition the Wright-Fisher diffusion to eventually be lost.The transition density can be computed as

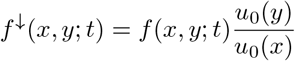

by using Bayes theorem. The diffusion conditioned on loss is characterized by its generator,

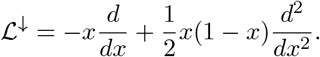

In an infinite sites model, in which mutations occur at the times of a Poisson process with rate *θ/*2 and then each drift according to the Wright-Fisher diffusion, a quasi-equilibrium distribution will be reached, known as the frequency spectrum. The frequency spectrum, *ϕ*(*x*), predicts the number of sites at frequency *x*, and can be written in terms of the speed measure and the probability of loss,

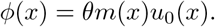

To proceed with calculating (4), note that the conditional probability of an allele being at frequency *y* in the ancient population given that it’s at frequency *x* in the modern population can be calculated

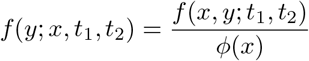

where *f* (*x, y*; *t*_1_, *t*_2_) is the joint probability of the allele frequencies in the modern and ancient populations and *ϕ*(*x*) is the frequency spectrum in the modern population.

Assuming that the ancestral population of the modern and ancient samples was at equilibrium, the joint distribution of allele frequencies can be computed by sampling alleles from the frequency spectrum of the ancestor and evolving them forward in time via the Wright-Fisher diffusion. This can be written mathematically as

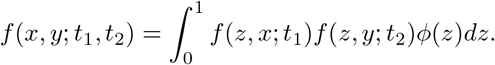

We now expand the frequency spectrum in terms of the speed measure and the probability of loss and use reversibility with respect to the speed measure to rewrite the equation,

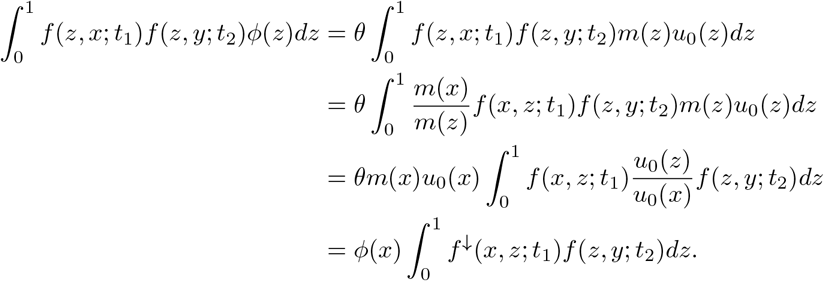

The third line follows by multiplying by *u*_0_(*x*)*/u*_0_(*x*) = 1. This equation has the inter-pretation of sampling an allele from the frequency spectrum in the modern population, then evolving it *backward* in time to the common ancestor, before evolving it *forward* in time to the ancient population. The interpretation of the diffusion conditioned on loss as evolving backward in time arises by considering the fact that alleles arose from unique mutations at some point in the past; hence, looking backward, alleles must eventually be lost at some point in the past.

To compute the expectation, we substitute this form for the joint probability into (4),

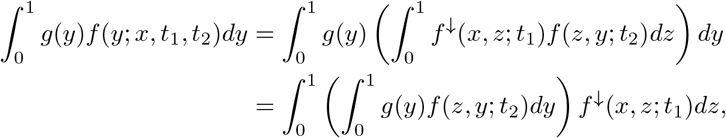

where the second line follows by rearranging terms and exchanging the order of integration. Note that this formula takes the form of nested expectations. Specifically,

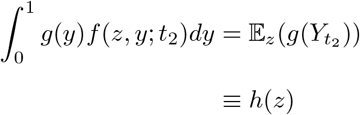

and

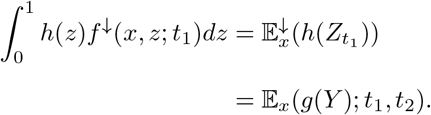

We now use (5) to note that

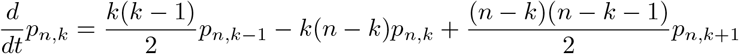

and

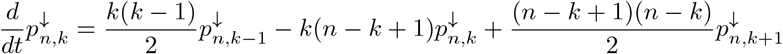

with obvious boundary conditions *p_n,k_*(0; *z*) = *z^k^*(1 *− z*)^*n−k*^ and 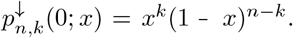

These systems of differential equations can be rewritten as matrix differential equations with coefficient matrices *Q* and *Q^↓^* respectively. Because they are linear, first order equations, they can be solved by matrix exponentiation. Because the expectation of a polynomial in the Wright-Fisher diffusion remains a polynomial, the nested expectations can be computed via matrix multiplication of the solutions to these differential equations, yielding the formula (2).

### 5.2 Sites covered exactly once have no information about drift in the ancient population

Consider a simplified model in which each site has exactly one read. When we have sequence from only a single individual, we have a set *l_a_* of sites where the single read is an ancestral allele and a set *l_d_* of sites where the single read is a derived allele. Thus, we can rewrite (3) as

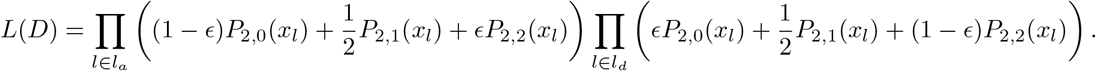

We can use formulas from Racimo et al. [2016] to compute *P*_2,*k*_(*x_l_*) for *k* ∈ {0, 1, 2},

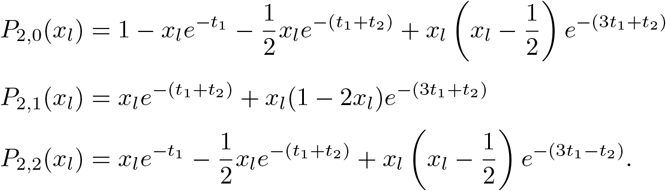

Note then that

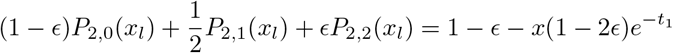

and

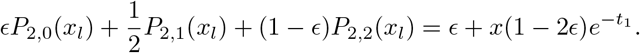

Neither of these formulas depend on *t*_2_; hence, there is no information about the drift time in the ancient population from data that is exactly 1x coverage.

### 6 Software Availability

Python implementations of the described methods are available at www.github.com/schraiber/con

## 7 Acknowledgments

We wish to thank Melinda Yang, Iain Mathieson, Pontus Skoglund, and Fernando Racimo for several discussions during the conception of this work that greatly improved its scope and rigor. We are grateful to Fernando Racimo and Benjamin Vernot for comments on early versions of this work that significantly improved its quality. We also appreciate comments on the preprint from Alex Kim and Aylwyn Scally. We would also like to thank Tamara Broderick for several stimulating discussions about clustering that ultimately didn’t result in any interesting application to ancient DNA.

